# Time-series genome-centric analysis unveils bacterial response to operational disturbance in activated sludge

**DOI:** 10.1101/565770

**Authors:** María Victoria Pérez, Leandro D. Guerrero, Esteban Orellana, Eva L. Figuerola, Leonardo Erijman

## Abstract

Understanding ecosystem response to disturbances and identifying the most critical traits for the maintenance of ecosystem functioning are important goals for microbial community ecology. In this study, we used 16S rRNA amplicon sequencing and metagenomics to investigate the assembly of bacterial populations in a full-scale municipal activated sludge wastewater treatment plant over a period of three years, including a period of nine month of disturbance, characterized by short-term plant shutdowns. Following the reconstruction of 173 metagenome-assembled genomes, we assessed the functional potential, the number of rRNA gene operons and the *in situ* growth rate of microorganisms present throughout the time series. Operational disturbances caused a significant decrease in bacteria with a single copy of the ribosomal RNA (*rrn*) operon. Despite only moderate differences in resource availability, replication rates were distributed uniformly throughout time, with no differences between disturbed and stable periods. We suggest that the length of the growth lag phase, rather than the growth rate, as the primary driver of selection under disturbed conditions. Thus, the system could maintain its function in the face of disturbance by recruiting bacteria with the capacity to rapidly resume growth under unsteady operating conditions.

**IMPORTANCE:** In this work we investigated the response of microbial communities to disturbances in a full-scale activated sludge wastewater treatment plant over a time-scale that included periods of stability and disturbance. We performed a genome-wide analysis, which allowed us the direct estimation of specific cellular traits, including the rRNA operon copy number and the in situ growth rate of bacteria. This work builds upon recent efforts to incorporate growth efficiency for the understanding of the physiological and ecological processes shaping microbial communities in nature. We found evidence that would suggest that activated sludge could maintain its function in the face of disturbance by recruiting bacteria with the capacity to rapidly resume growth under unsteady operating conditions. This paper provides relevant insights into wastewater treatment process, and may also reveal a key role for growth traits in the adaptive response of bacteria to unsteady environmental conditions.

## INTRODUCTION

Understanding the drivers of community structure is important for developing predictive models and for guiding engineering and management practice of microbial community ecosystems (1–4). Activated sludge is the most widely used process for biological wastewater treatment worldwide (5). It involves highly diverse biomass aggregated into flocs, which can be separated from treated wastewater by gravity settling. The heterogeneous structure of activated sludge flocs is a result of physical, chemical, and biological processes. These include (a) physicochemical interactions between microorganisms, exopolymeric substances (EPS), inorganic particles and di-and trivalent cations, (b) biological interactions among diverse microorganisms within the community, and (c) physiological processes for nutrient uptake and resource allocation between biological activities in bacterial cells (6). The selection of microbial populations in these ecosystems is driven primarily by environmental and operational pressures (7). A great deal of knowledge has been achieved during the last two decades from research exploring how bacterial community composition is affected by process configuration, solid retention time (SRT), temperature, redox conditions, wastewater composition, pH, and other factors (8–16). Recent progress in high-throughput sequencing technology has facilitated a detailed characterization of the composition of microbial communities in a large number of wastewater treatment systems worldwide (12, 17–19). However, the ability to produce a precise ecological description of activated sludge based on the abundance of the species present is still elusive.

Typically, wastewater treatment plants (WWTP) are able to perform reliably under fluctuating conditions. Given that the stability of microbial communities is affected by disturbance events, it is important to understand how ecosystems respond to disturbance and which traits are the most critical for the maintenance of functioning. To date few studies have addressed the response of microbial communities to disturbance in full-scale wastewater treatment systems (20). Vuono et al. observed that a sudden decrease in SRT from 12 to 3 days, prompted a shift in community structure, favoring members of *Burkholderiales* and *Sphingobacteriales*, two taxa defined as fast-growing organisms that are adapted for high resource availability (“r-strategists”) (11).

Identifying traits that explain the ability of species to respond to disturbances may also allow for greater insight into the microbial ecology of the process beyond the description of community composition. Activated sludge systems operate at loading rates typically in the range of 0.05 to 0.4 kg organic matter per kg dry biomass and per day, and can be therefore considered oligotrophic environments (21). That means that under stable operation, bacteria must adapt to grow under conditions of carbon limitation (11, 22). Indeed, it has been inferred on the basis of the average number of *rrn* operons, that activated sludge are dominated by bacterial populations that are near carrying capacity and make efficient use of available growth-limiting resources (11, 23). Thus, ecological selection might also be acting at the level of metabolic potential of bacterial populations. This hypothesis could be tested directly by examining signatures in the genomes of the bacterial community. Metagenomics offers the possibility to capture the genomic complexity of bacterial communities at high resolution. In particular, genome-centric approaches have the additional advantage that specific traits can be analyzed in the context of the other functional properties of the microorganisms (24).

In this study, we used metagenomics and 16S rRNA sequencing to investigate bacterial population and functional dynamics in a full-scale municipal activated sludge wastewater treatment plant over a period of three years, which comprised distinct operational process conditions. Similar studies that have investigated the importance of functional traits for community assembly processes applying the PICRUSt approach (25) to predict functional profiles of microbial communities using 16S rRNA gene sequences, and have estimated the number of rRNA gene operons using the Ribosomal RNA Database (11, 26). Our analysis differs importantly from those previous studies in that we accomplished genome reconstruction to assess the functional profile, and we estimated the number of rRNA gene operons and the *in situ* growth rate of assembled genomes (27, 28). We first used 16S rRNA amplicon sequencing data to identify patterns of co-occurrence between taxa over the three-years-time series, and compared the distribution of community composition with operational and performance metadata, thereby establishing a consensus for periods of disturbance and high stability. Secondly, we used metagenomic data to identify differences in functional profiles between contrasting operational periods. Finally, we estimated the number of rRNA gene operons and the *in situ* growth rate of reconstructed genomes across the time series that comprised distinctly perturbed and stable periods, to test the hypothesis that shifts in bacterial community structure caused by process disturbances reflected differences in functional characteristics of microorganisms, including their growth traits.

## RESULTS

### WWTP performance

The main characteristics and average performance data of the municipal full-scale activated sludge plant are given in Table S1. The sewage treatment plant was sampled during the course of a planned capacity expansion, which gave rise to a singular period, which lasted approximately nine months, characterized by repeated disturbances, caused mainly by short-term plant shutdown, each time for several hours. Features of this period were interruption of wastewater feed supply, fluctuations in mixed liquor suspended solids (MLSS) concentration, and/or eventual periods of very low dissolved oxygen concentration, resulting in higher effluent chemical oxygen demand (COD) (Fig. 1A). After a new operative module was gradually put into operation, there was a moderate decrease in food to microorganism ratio (F/M) (Fig. 1B), which ultimately led to a late period of nine months of very good performance, which included full nitrification (Fig. S1). The SRT, ranging between 4 and 7 days, changed in the opposite direction of F/M.

**Fig. 1:**
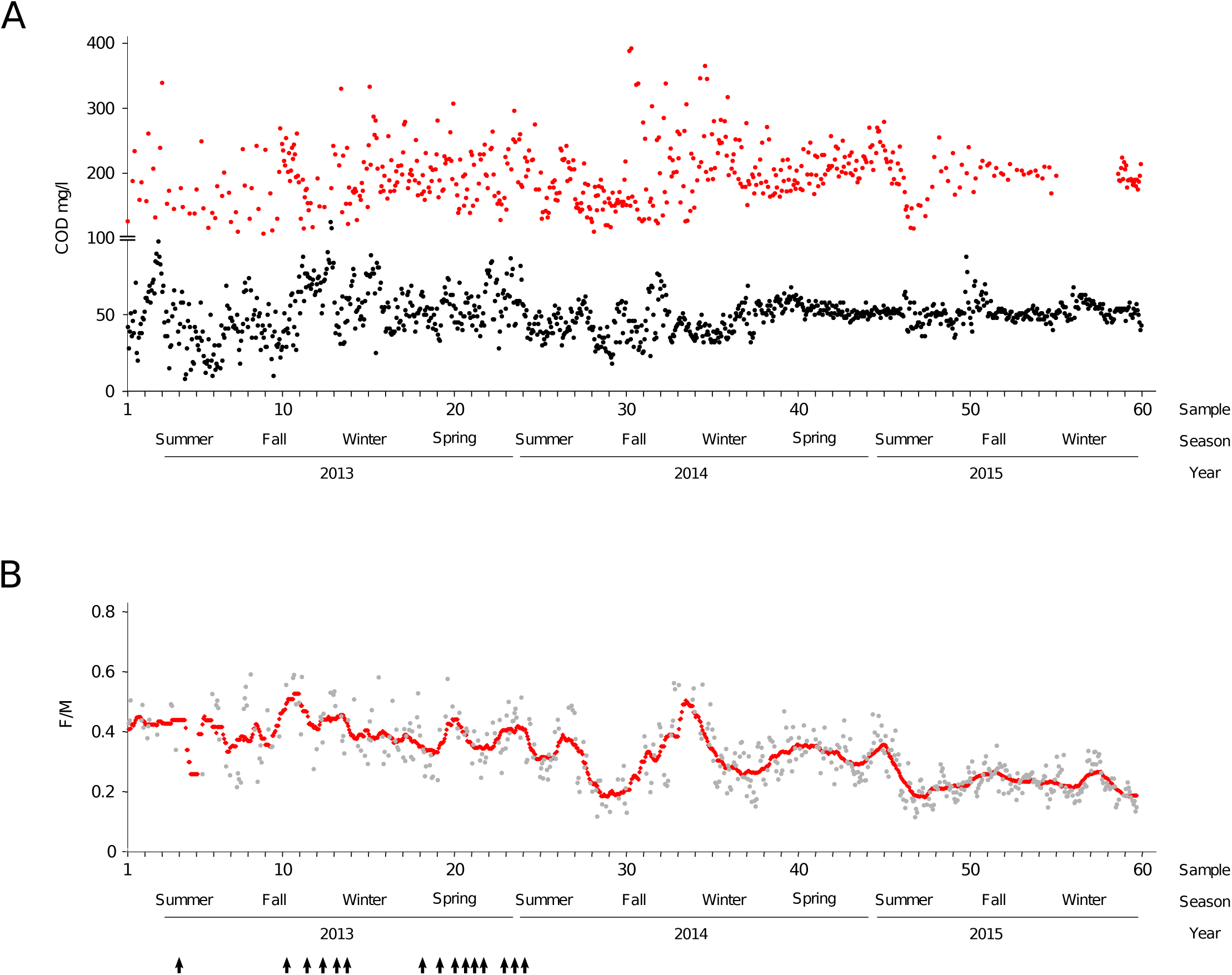
Functional performance and operating conditions of the full-scale activated sludge over three years of sampling. A) Influent (red dots) and effluent (black dots) chemical oxygen demand concentration. B) Food: Microorganism (F:M) ratio. Red dots indicate the F/M calculated using data based on a 28-day moving average (89). Black arrows indicate disturbance events.

### Phylogenetic analysis of activated sludge bacterial community structure and dynamics

Initial analysis of the three years-time-series was based on sequences of 16S rRNA gene amplicons, which were assigned to 1002 OTUs (97% similarity) with relative abundance higher than 0.01%. We performed LSA on the most abundant bacterial OTUs in order to explore co-occurrence patterns. The network analysis grouped bacterial OTUs into two main clusters, which were negatively correlated with one another (Fig. 2A). OTUs corresponding to one of the main clusters belong mostly to phyla *Bacteroidetes* and Proteobacteria, whereas phyla Acidobacteria, Actinobacteria, Patescibacteria, Chloroflexi, Planctomycetes and Nitrospira, were mostly represented in the other major cluster. A plot of the temporal distribution of bacteria belonging to the two main clusters of the network shows a striking correspondence to each of the two operationally distinct periods of disturbance and stability (Fig. 2B).

**Fig. 2:**
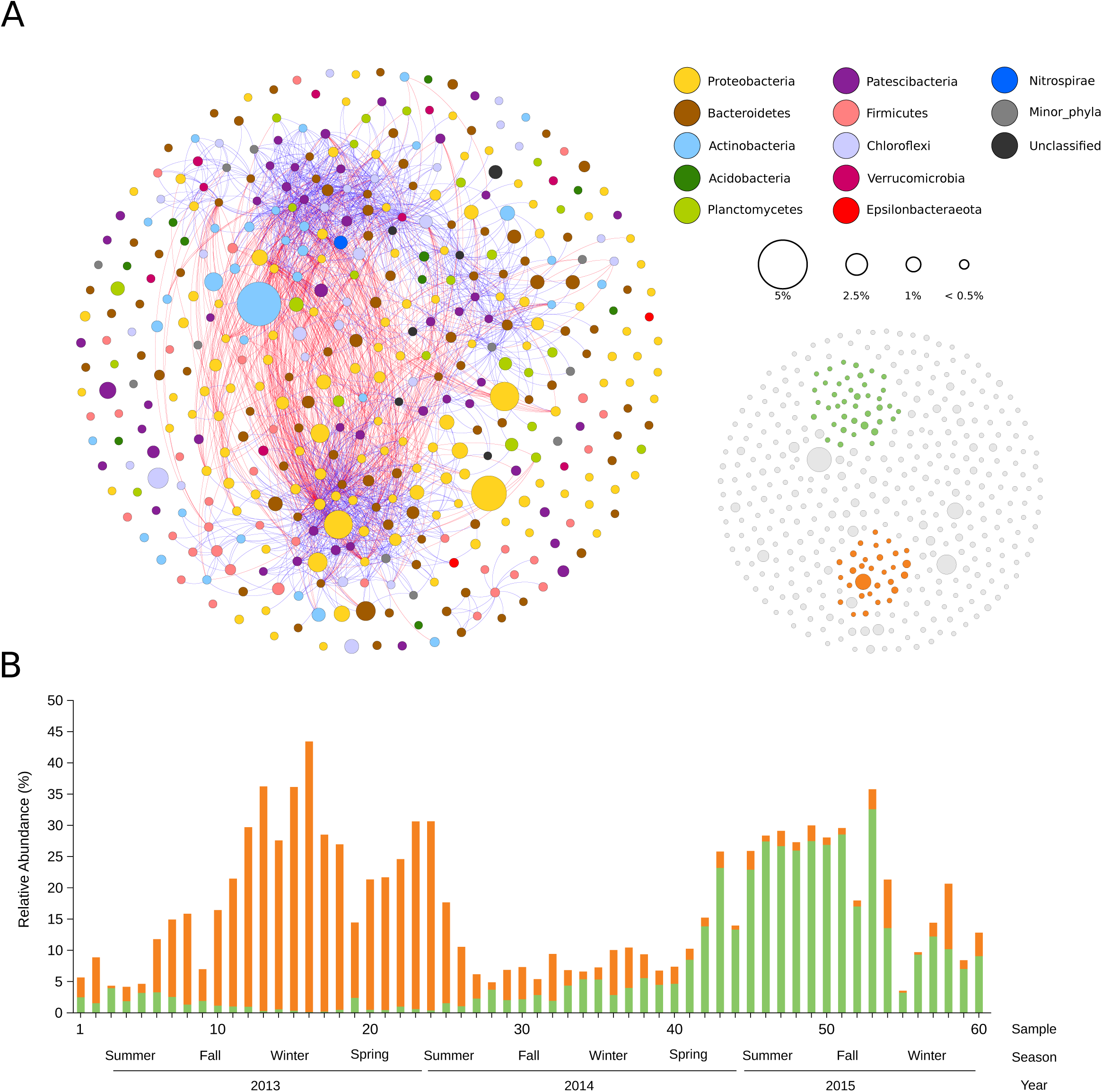
A) Network of co-occurring bacterial OTUs, based on local similarity analysis (LSA). Nodes (OTUs) are colored by phylum. The size of each node is proportional to the OTU relative abundance across the 60 samples. Blue and red edges represent positive and negative significant correlations, respectively (correlation > |0.6|; P-value < 0.01). Nodes that have no connections are distributed randomly. Inset: The application of a k-core filter = 10 (considering only positive correlations) highlights two clusters (nodes in each cluster are denoted in green and orange). B) Time series abundance of OTUs belonging to the two major clusters of the network shown in panel A. Orange and green bars are the sum of the relative abundance of nodes (OTUs) in each cluster.

Further taxonomic analysis was performed on drafts genomes assembled from metagenomes. We were able to reconstruct 173 good quality MAGs, taking advantage on the differences in contig coverage across the large number of metagenomic datasets (n=60). Table S2 details the characteristics of all MAGs, including their taxonomic affiliation, the length, number of contigs, degree of completeness and contamination. Similar to the results obtained for amplicons, network analysis of MAGs also showed two distinguishable clusters (Fig S2).

Given that rRNA genes of microbial communities are in general not efficiently recovered using *de novo* assemblers, we used an iterative mapping method, based on the expectation maximization algorithm to reconstruct full-length SSU sequences from raw reads (29). In order to evaluate whether the population sets obtained from the reconstruction of genomes in the metagenomes was a fair representation of the “true” bacterial community structure, we compared the average bacterial community structures determined from: 1) the taxonomic classification of 16S rRNA gene amplicons (Amp16S), 2) the taxonomic annotation of 16S rRNA gene sequences assembled from the metagenomes (MA16S), and 3) the taxonomic placement of metagenome assembled genomes (MAGs) (Fig. S3).

As expected, the distribution of taxa at higher levels were very similar, albeit not identical. The higher sequencing depth afforded by amplicon-sequencing allowed to solve higher proportion of minor phyla. Additionally, incomplete phylogenetic placement of amplicons resulted in a small proportion of unclassified taxa. Other differences could be explained by the different sources of amplification biases occurring in PCR-based methods, the bias in the estimation of taxa relative abundances due to differences in *rrn* operon copy number, and the incomplete reconstruction of the total community members as MAGs. Despite these differences, the three methods showed a highly satisfactory agreement. Therefore, we conclude that the set of genomes assembled from the metagenomes adequately represent the bacterial communities present in the system, confirming that our sequencing effort provided sufficient resolution to obtain a comprehensive insight in the prokaryotic microbial community of the system.

### Metagenomic analysis

The distribution of taxa across the time series was investigated with MAGs as input to construct 5×5 self-organizing maps (SOM) (Fig S4). SOM grouped MAGs into two distant superclusters (colored in green and orange in Fig. 3A), and a third supercluster (pink-colored) that includes all other sampling dates. In order to identify differences in functional composition, an analogous SOM analysis was applied to the whole metagenome (Fig 3B). Relative abundance of annotated genes (KEGG modules) in metagenomic contigs resulted in a similar clustering pattern, with two distant groups of samples (also colored in green and orange), and two additional periods (pink and purple) that include the remaining samples. Comparison between functional-based and genome-based SOM analyses enabled us to establish two consensus periods, characterized by their close correspondence to operational periods of disturbance (samples 10 to 25) and stability (samples 42 to 58) (Fig. 3C).

**Fig. 3:**
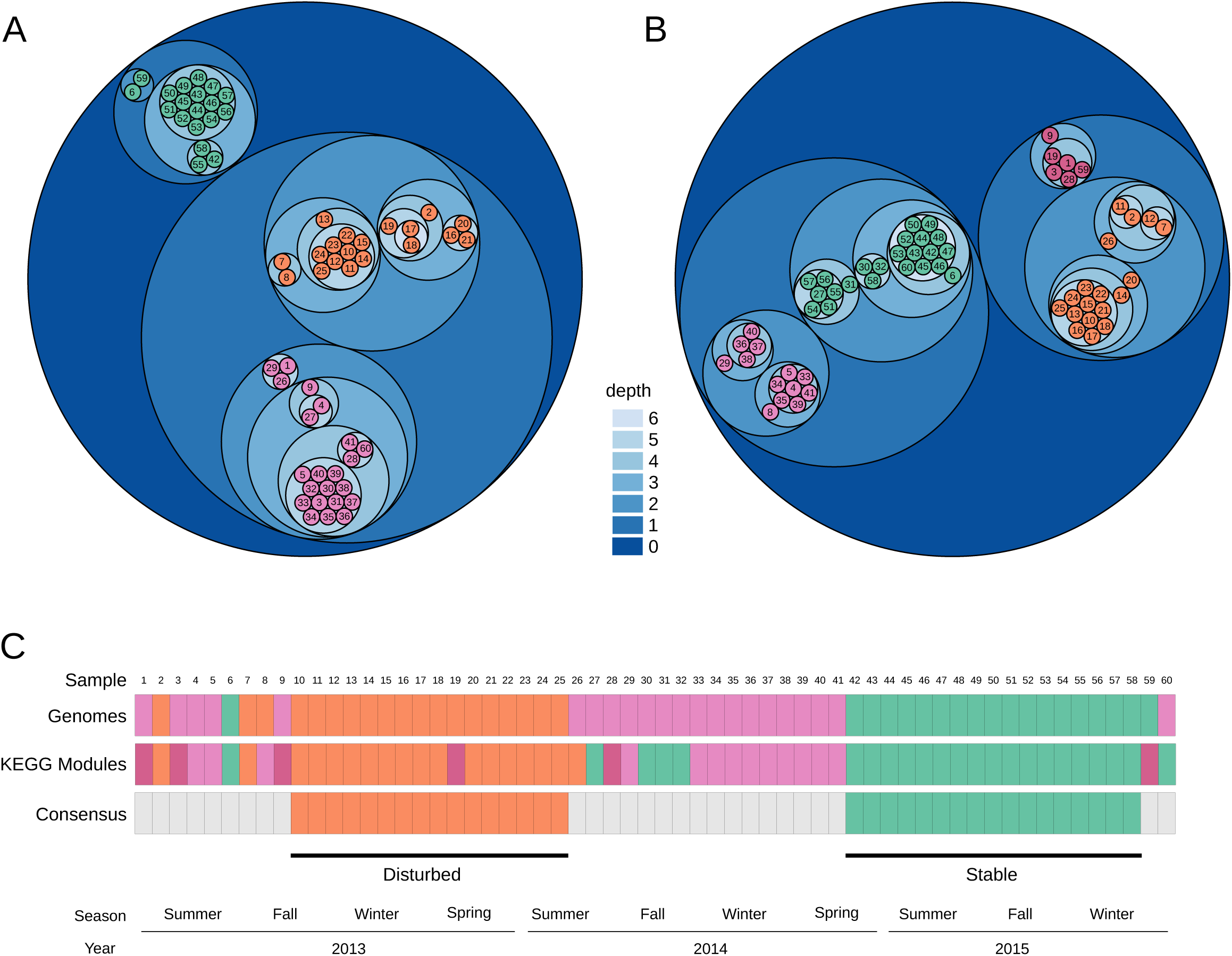
Self-organizing map (SOM) analysis. Circle packing representation of the tree derived from the 5×5 SOM clustering results (Fig S4). Cluster nodes are visualized as nested circles, with the node levels depicted by a blue gradient. Numbers in childless nodes are those of the time-series samples. Samples grouped into superclusters were classified using Ward’s minimum variance method, and are displayed in orange, green, pink and purple. A) SOM clustering based on MAGs abundance and B) SOM clustering based on KEGG module abundance. C) Overlap between samples in superclusters from A and B were used to define consensus samples for disturbed (orange) and stable (green) periods.

Comparison of functional profiles between metagenomes from disturbed and stable periods disclosed a number of pathways that had significant differences in relative abundance (Fig. S5). The disturbed period was significantly enriched in genomic features such as secretion system, two component systems, transporters and energy metabolism. On the other hand, the stable period contained a significantly higher proportion of genes in the categories of metabolism and genetic information processing and genes coding for biosynthesis of secondary metabolites and genes (Fig. S5).

Statistical comparisons were also used to discriminate MAGs according to differences in their abundance between disturbed and stable periods (Fig. S6). In agreement with the 16S-based results, the abundance of Proteobacteria increased during the disturbed period. Nevertheless, most phyla have representatives in both disturbed and stable periods. Likewise, a genome-wide comparison of MAGs, based on predicted metabolic capacities (KEGG modules), indicated that phylogenetically-related MAGs, sharing a large number of annotated genes, split into disturbed and stable periods (Fig. S7).

### Bacterial community structure and growth traits

We searched for traits encoded in bacterial genomes that could account for the differences observed in bacterial composition between disturbed and stable periods. Firstly, we calculated the ribosomal RNA (*rrn*) operon copy numbers. We applied an approach for quantifying *rrn* copy numbers of MAGs, which is independent on the presence of *rrn* operons in the assembled genomes. Matching MAGs to their corresponding 16S rRNA was initially accomplished by Pearson correlation of the coverage of the MAG and the coverage of the 16S rRNA gene sequences reconstructed independently (MA16S) along the entire time series. The matching was considered robust if the correlation was *r* > 0.7 (p< 0.001) and the taxonomic classifications based on the 16S rRNA sequence and on the genome phylogeny were coincident (Table S3). Further support was obtained from BLAST and/or pair-end reads connecting the 16S rRNA gene sequences assembled from the metagenomes (MA16S) with contigs in the MAG (see M&M). The filtering procedure was successfully applied to 49 MAGs, for which we could unequivocally matched the reconstructed 16S rRNA gene with the assembled genome. The number of ribosomal RNA operons (*rrn*) in each MAG was subsequently calculated from the relationship between the coverage of the MA16S and the average coverage of the rest of the genome (Fig S8 and Table S3). Results obtained using this approach was further checked by comparing the estimated *rrn* copy number against the copy number retrieved from *rrndb*, the ribosomal RNA operon copy number database (30) (Table S3). Fig. 4A shows that the proportion of genomes with a single copy of *rrn* operon was highest during the period of stable operation and decreased to a minimum during the period of plant disturbance. Differences in the abundance-weighted mean of per-MAG rRNA operon copy number between both contrasting periods were significant at P <0.001 (Fig. 4B). An almost identical result was obtained when the *rrn* copy number was calculated using the contigs directly from metagenomes, as the ratio of the coverage of total rRNA SSU genes (MA16S) to the average coverage of total genomes, determined from the coverage of a set of single-copy marker genes (Fig 4C and 4D).

**Fig. 4:**
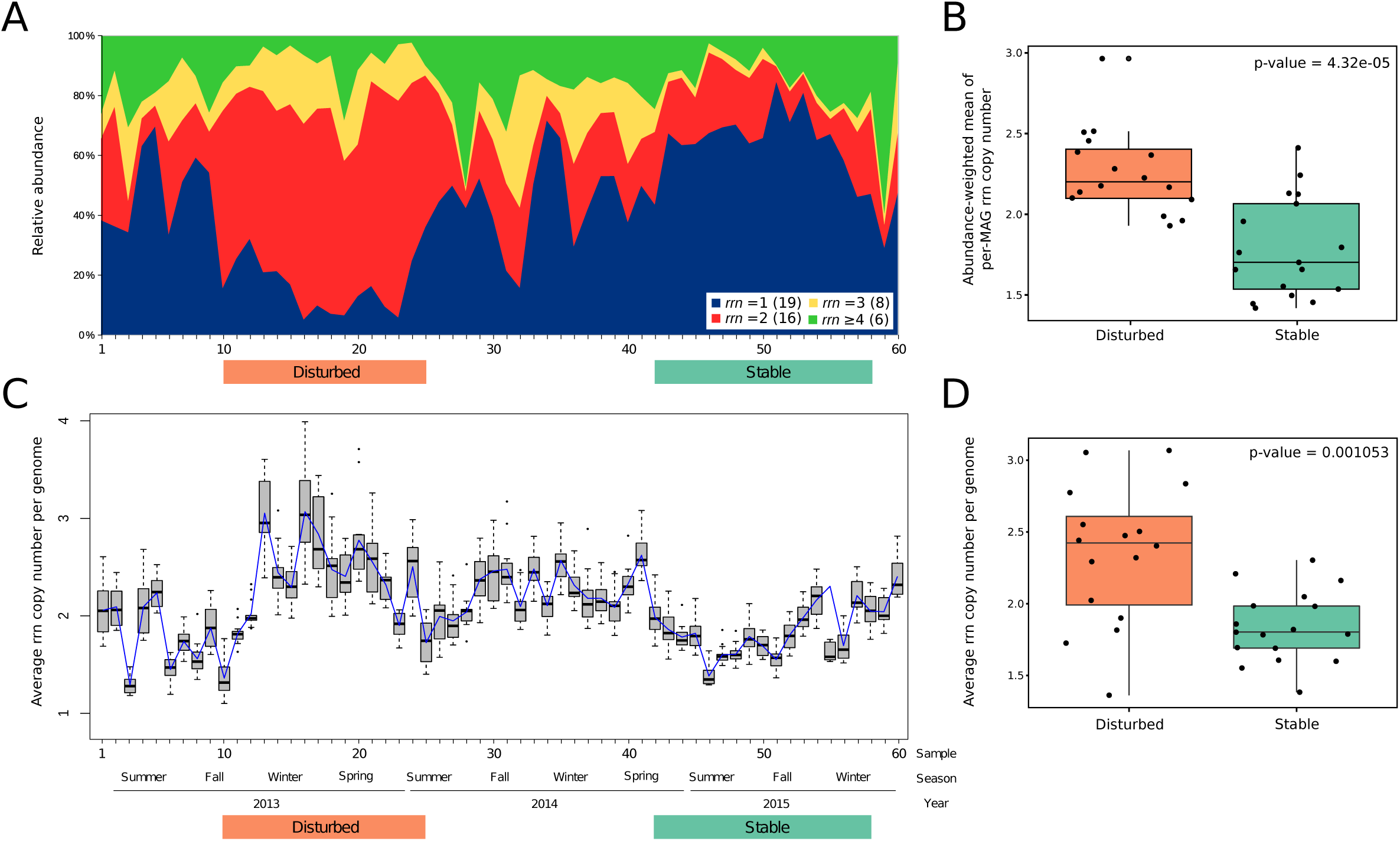
A) Relative abundance of metagenome assembled genomes across the three-year time-series, color-coded according to the number of copies of the *rrn* operon. Colored bars delimit samples corresponding to disturbed (orange) and stable (green) consensus periods defined in Fig 3. In parenthesis are the number of MAGs in each category. B) Boxplot of abundance-weighted mean of per-MAG *rrn* copy number, calculated from all MAGs present in the periods marked as disturbed (orange) and stable (green). C) Average copy number of ribosomal RNA operons (*rrn*) per genome, calculated as the ratio of the coverage of total rRNA SSU genes to the total coverage of single copy universal marker genes in metagenomes’ contigs. The boxplot was generated using 13 single-copy marker genes. The blue line is the median. D) Boxplot of the average *rrn* copy number per genome in the periods marked as disturbed (orange) and stable (green).

Because *rrn* copy number is directly related to maximum growth rate in bacteria (31), we reasoned that differences in *rrn* copy number could be reflected by changes in bacterial growth rate between the disturbed and stable periods. To test this idea, replication rates were estimated across the time series using the index of replication (iRep). The index was estimated for 113 points in the 35 genomes that were ≥ 75% complete and had coverage higher than 5 in each individual sample (Fig. 5A). In contrast to our prediction, iRep values were distributed uniformly throughout time, without differences between disturbed and stable periods, and with no relation to any other measured parameter, including F/M. The null hypothesis that the distribution of iRep values are indistinguishable from a normal distribution centered at 1.57 with a standard deviation of 0.14 could not be rejected (p=0.39). This was further demonstrated by a very good fit between the quantiles of the iRep data and those of a normal distribution (Fig. 5B and Fig. 5C). Also, we did not find a significant correlation between the copy number of *rrn* operons and iRep values (n=33, Spearman’s correlation r=0.31, p=0.082). Based on these observations, we conclude that bacteria present in disturbed and stable periods are characterized by differences in *rrn* copy number but not by growth rate.

**Fig. 5:**
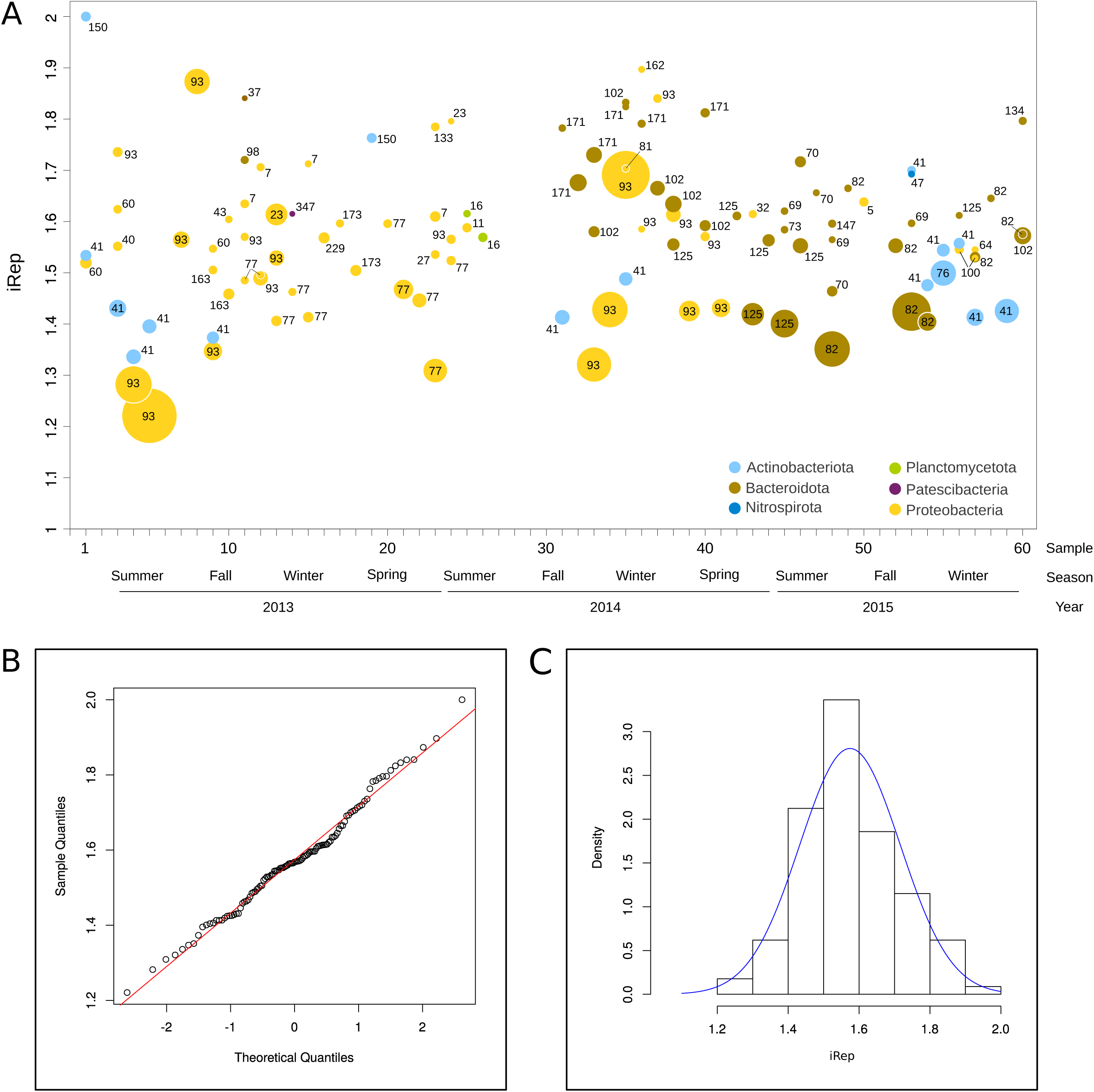
A) Replication index (iRep) calculated from the difference of coverage between the regions of the genome close to the single replication origin and those of the rest of the metagenome assembled genomes. Values of iRep were calculated for 35 MAGs across 60 samples. Phyla are distinguished by colors. The size of the circles is proportional to the MAG coverage. Numbers denote MAGs according to Table S2. B) QQ plot of iRep values versus normal distribution. C) Histogram of iRep values. Solid line in blue represents the normal distribution centered at 1.57 with a standard deviation of 0.14.

## DISCUSSION

A genome-centric approach using time-series metagenomic data of activated sludge samples allowed us the characterization of community response to disturbance. Our results revealed clear differences between disturbed and stable operational periods as regards to bacterial ribosomal RNA (*rrn*) operon copy numbers. On the other hand these differences did not reflect in corresponding changes in replication rates. We interpret these results to suggest that the length of the growth lag phase, rather than the growth rate, could be the primary driver of bacterial selection under disturbed conditions.

A stable activated sludge process is a system that produces effluent with very low oxygen demand, thus representing an environment characterized by a constant but slow flux of resources, meaning that bacteria are fed at near-starvation rate (21). Bacteria have developed a wide range of growth and survival strategies to maximize their reproductive success in such environments. The trade-off between resource use efficiency and growth rate explains the dominance of bacteria harboring low *rrn* operon copy number as the result of a negative selection for fast growing bacteria (32), and the adaptation of bacteria with higher growth efficiency in low resource environments (31, 33). In agreement with these theoretical framework, our results showed that bacteria with low copy number of *rrn* operons dominated the activated sludge community during periods of optimal performance. Similar results were obtained by Vuono and colleagues, who manipulated the SRT of a full-scale wastewater treatment plant, and showed that phyla abundant at high and moderate SRT, such as Planctomycetes, Chloroflexi, Acidobacteria and Nitrospira, were outcompeted by fast-growing microorganisms that are adapted for high resource utilization, such as Proteobacteria upon a shift to very low SRT operation (11). A positive linear correlation between resource availability and the average rRNA operon copy number of microbial populations was also observed in anaerobic digesters (34). Comparable trade-offs between bacterial ecological strategies based on *rrn* copy number distribution were uncovered in microbial succession in several habitats (35), including soil microcosms (36), and stream biofilms (37). In all these previous studies, the observations were rationalized using the r-K selection model, based on life-history strategies. Classification of organisms into r-strategists and K-strategists provides a conceptual basis for making specific predictions about the ecological attributes and niche preferences of organisms (38), including microorganisms (23, 39). In simplified terms, *r*-strategists dominate when resources are abundant, due to their ability to grow exponentially using the available resources. K -selected species, on the other hand, are adapted to compete when populations are near carrying capacity and resources are in short supply (38). However, growth traits of an organism cannot be directly inferred from taxa-based classifications, especially when oligotrophic microorganisms may be r-strategists, whereas copiotrophic microorganisms may be K-strategists (40). As a case in point, we found in this study members of the same family that contain different *rrn* copy numbers and are abundant during periods of contrasting stability (see eg *Polyangiaceae* in Fig S7), confirming the heterogeneity in life-history strategies among phylogenetically related taxa (41). The fact that life-history traits of microorganisms present throughout disturbed and stable periods were estimated directly using the experimental data is a strength of this paper. The possibility of inferring genomic traits from high quality drafts genomes extracted from metagenomes highlights the value of organism-centric approach in metagenomics (24).

Based on the generalized relationship between *rrn* copy number and bacterial growth rate (31, 42), we anticipated that differences in *rrn* copy number would mirror differences in bacterial growth rate between the disturbed and stable periods. Evaluating bacterial growth rate of specific lineages could be performed using a mass balance, if the numbers of bacteria entering and leaving the system are known (17). Alternatively, growth rates can be probed in metagenomic samples from the differences in read coverage across bacterial genomes (43). Indeed, the algorithm iRep is already an established tool for calculating the inferred *in situ* replication rates of bacteria in microbial communities (27, 44–46).

We found in this study that, even though bacteria with higher *rrn* copy number dominated the disturbed period, estimated replication index values were distributed uniformly throughout time, and no differences were observed between disturbed and stable periods, nor had any relation to SRT. We note that even though the large number of data points gives us high confidence in the reliability of our data, we cannot prove that MAGs used for iRep calculation were a truly unbiased representation of the community. Therefore, actual unseen differences in growth rate cannot be entirely ruled out.

Yet bacterial growth traits that contribute to fitness include not only the rate of exponential growth, but also the length of the lag phase (47). Thus, we infer from our results that shorter lags could provide bacteria with high *rrn* copy number the competitive advantage to rapidly adjust growth under unsteady operating conditions. Recent experiments support our proposition. It has been shown that the length of the growth lag phase declines with the number of ribosomal operons (32, 42, 48), and that the competitive fitness difference between *E. coli* strains with different numbers of *rrn* operons depends on the dynamics of nutrient availability, being most pronounced during the lag phase (49). Similarly, computational models and benchtop experiments of pairwise competition between *E. coli* mutants of the essential enzyme adenylate kinase (Adk), demonstrated that strains with shorter lag time exhibit higher competitive advantage, especially under nutrient-limiting conditions (50).

We expect that this hypothesis could be confirmed in future research. Direct measurement of the lag phase, using methods such as total viable count, optical density measurements or even qPCR with primers targeting specific bacterial populations, is not readily attainable in the context of complex microbial communities such as activated sludge. On the other hand, accurate prediction of the lag phase using computational approaches are very difficult to obtain even for simple model sytems (51).

One of the objectives of shotgun analysis is to uncover possible functional genomic signatures that advance our understanding of the mechanisms that allowed bacteria to adapt to contrasting regimes of stability and disturbance in activated sludge, and how they were reflected in their genomes. The molecular basis of bacterial response to the environment are still poorly understood. Existing studies are based on the exploration of genomes of marine microorganisms (52, 53). In general terms, we note that functional capabilities did not predict which bacterial populations will dominate periods of disturbance or stability. On the contrary, MAGs belonging to each of the contrasting operational periods were not necessarily functionally related. Analysis of MAGs belonging to disturbed and stable periods, based on the presence or absence of metabolic pathways inferred from KEGG database, indicated that MAGs clustered by phylogeny, rather than by the operational regime to which they belong (Fig S7). The failure to identify a prototypical genome content that allows clear distinctions between copiotrophic or oligotrophic bacteria was discussed previously by (31). Yet we observed differences at the genomic level that are consistent with recently proposed models, which predict that competitive advantage of bacteria under changing environmental condition is achieved by reserve capacity of ribosomes and transporters. Accordingly, sub-saturation allows bacteria to rapidly adapt in fluctuating environments by producing new ribosomes and high affinity transporters for optimal growth (54, 55). We also observed that the disturbed period was significantly enriched in secretion system, which are used to deliver a variety of different proteins, including bacterial toxins and degradative enzymes such as proteases and lipases (56). Two-component systems, used by bacteria to detect changes in their environment (57) were also overrepresented in the disturbed period. On the other hand, the stable period contained a significantly higher proportion of genes coding for increased production of secondary metabolites, which are characteristics of organisms with slow but efficient lifestyle (52).

It may be argued that the nature and frequency of the disturbances occurring on our field-scale experiment were relatively ill-defined. In general, disturbances can be difficult to define, especially for observational studies (58, 59). Here, we concur with the definition of disturbance, recently put forward by Cante for use in microbial ecology: “A discrete, unpredictable event that causes direct removal of living biomass, thereby altering community structure” (60). Of interest is that this definition refers to discrete and unpredictable events, and that it does not include the notion of disturbance as a rare or relatively infrequent event. In our experiment, disturbance is a random factor that encompassed several events, although the length of the interval between disturbances exceeded the short generation times of microorganisms.

At first sight, the suggestion that growth rate was approximately invariant over a range of F/M, SRT, temperature and even operational conditions may seem unsettling to the wastewater practitioner. However, it is not entirely unexpected, since many authors have found that the yield coefficient Y_H_ of heterotrophic biomass is independent of temperature (61). and there is experimental evidence that abundant taxa occur at a wide range of sludge ages (62). Furthermore, it has long been recognized that changes in solid retention time had only a slight effect on the sludge production coefficient (63). Thus, specific growth rate is ultimately determined by the low concentration of readily biodegradable substrate in the reactors.

In summary, our results show that disturbances, defined by short-term interruption of wastewater feed supply, fluctuations in MLSS concentration, and/or eventual periods of very low dissolved oxygen concentration, increase the relative proportion of bacteria with higher rRNA operon copy number. Given that the *rrn* operon copy number is considered to reflect ecological strategies in bacteria by influencing their growth traits, we suggest that the length of the growth lag is of primary importance for the capacity of bacteria to thrive under disturbance. It has been shown that r-strategists are more resilient to disturbances than K-strategists (40, 65–67). Our results give experimental support to the prediction that communities with frequent disturbance regimes will be dominated by more resilient communities (40). We propose that the system has the capacity to maintain its function in the face of disturbance (ecological resilience) through the selection of bacteria that able to return rapidly to their equilibrium or steady–state condition (engineering resilience). Future experiments examining the time course of microbial composition after disturbance using shorter sampling intervals could provide a direct experimental test for the hypothesis put forward in this study, i.e. that a shorter lag phase provided bacteria an advantageous trait in a disturbed environment.

## MATERIALS AND METHODS

### Sample collection

Samples were obtained from a full-scale municipal WWTP, located in the metropolitan area of Buenos Aires (Argentina), which provides preliminary, primary and secondary treatment to remove organic matter and suspended solids from sewage for a population of 600 000 residents. The WWTP has a modular design, with a capacity of each module to treat 78 000 m^3^/day. Biologic treatment system of each module consists in conventional activated sludge and includes four aeration chambers and four secondary clarifiers. At the time sampling started, only one module was under operation. Works for wastewater treatment plant upgrade started shortly after the beginning of the sampling period. The transition period was marked by events of operational anomalies, especially caused by temporary (several hours) shutdown of one or more processes of the WWTP, necessary for civil engineering works. Start-up of the second module was carried out gradually, according to the increase in total influent flow. Throughout the entire sampling period, the plant achieved satisfactory biochemical oxygen demand (BOD) removal, producing an effluent complying with the local effluent-quality regulation.

A total of 60 samples of activated sludge were collected bi-weekly over a period of three years (beginning on November 2012) from one of the aeration basins. Samples were transported within 2 h from the sewage plant to the laboratory, and stored at −70°C until DNA extraction.

### DNA extraction and sequencing

Total DNA from sludge samples was isolated by a direct lysis procedure involving physical disruption of cells and a CTAB method as described in (68). The 60 DNA samples extracted from sludge were sent to INDEAR, Rosario, Argentina, for Nextera DNA library preparation and sequencing. A rapid run sequencing on two lanes was performed in a HiSeq 1500 Illumina, generating paired-end (PE) reads of 250 bp. The same DNA samples were also sent to Macrogen Inc., Korea, for 16S ribosomal RNA gene amplicon sequencing. Amplicons of the V3-V4 region were sequenced using Illumina MiSeq, generating PE reads of 300 pb, using the primers b341F (5’-CCTACGGGNGGCWGCAG-3’) and Bakt805R (5’-GACTACHVGGGTATCTAATCC-3’) (69). Sequencing data are available at NCBI BioProject under accession PRJNA484416

### 16S rRNA amplicon sequencing analysis

Amplicon sequencing of the activated sludge samples resulted in 7 293 800 PE reads. After sequence quality controls, raw reads were filtered with Trimmomatic (70) using recommended quality parameters for PE reads and removing reads below 200 bases. A total of 4 581 297 PE reads passed the quality filters. USEARCH v8.1.1861 (71) was used to join the paired sequences, considering a minimum overlap of 32 bp. USEARCH pipeline was also used to filter merged sequences and define operational taxonomic units (OTUs) at 97% similarity. For diversity analysis, samples were rarified to the lowest number of sequences (11 810). Sequences were classified with v. Silva database v. 132 (72), using 88% minimum identity with the query sequence.

### Metagenome assembly

Metagenomic sequencing of the activated sludge samples resulted in 328 million PE reads (average of 5.5 million per sample). PE reads (2 × 250 bp) were filtered and trimmed to remove ambiguous bases (N) and ensured a minimum average quality value (Q) of 30. After quality filtering, approximately 35% of reads were removed and the remaining reads were used for the assembly. The filtered reads of the 60 samples were combined into a single file and assembled using MEGAHIT (73), with multiples k-mer length (95 to 99) and default parameters, generating 409 381 contigs longer than 1500 bp. In order to assemble individual genomes from the contigs pool, contigs were binned with MetaBAT (74), using tetranucleotide frequency and coverage information in each of the 60 samples. Binning was manually refined using Cytoscape (75) to visualize the following three criteria: a) GC percentages, b) tetranucleotide frequency distribution c) abundance profiles along the 60 samples. Manually binning of scaffolds consisted in clustering scaffolds (i.e. considered a bin) when the composite sequences contained ± 5% of average %GC and all scaffolds were interconnected according to Pearson’s significant positive correlations (based on tetranucleotide frequency distribution and abundance profiles). As a final step, the reads mapping the contigs from each bin were extracted and reassembled using MEGAHIT as described above. CheckM (76) was used to estimate the quality (completeness and contamination) of the metagenome-assembled genomes (MAGs).

To estimate the abundance per sample, filtered PE reads were mapped back to the contigs using Bowtie2 (77). The coverage was calculated using MetaBAT (jgi_summarize_bam_contig_depths script). Taxonomic assignment of bacterial genomes was done using the software toolkit GTDB-Tk v. 0.1.3 (78) and default parameters.

### Number of ribosomal RNA operons (rrn)

We applied an approach for quantifying *rrn* copy numbers in MAGs, which was independent on the presence of *rrn* operons in the assembled genomes. Reads corresponding to the 16S rRNA gene were identified from raw data of the 60 metagenomic samples using Metaxa2 (79) and assembled *de novo* using EMIRGE (29). Metagenome-assembled 16S rRNA genes (minimum 900 nt, called MA16S by analogy to MAG) were classified using the Silva database v. 132 (72), using 88% minimum identity with the query sequence. Filtered PE reads were mapped back to the MA16S with Bowtie2, and the coverage per sample was estimated using MetaBAT as described above. We applied several filtering criteria to ensure the most-accurate matching between MA16S and MAGs for use in downstream analyses. MAGs were initially matched to their corresponding 16S rRNA by Pearson correlation (r > 0.7, p< 0.001) between the coverage of the MAG and the coverage of MA16S along the entire time series. Secondly, taxonomic classifications based on the 16S rRNA sequence and on the genome phylogeny had to be concurrent at the highest possible taxonomic resolution. Further support was obtained by sequence match between assembled 16S rRNA genes and contigs in the MAG using BLAST (98% similarity; alignment length > 100 nt; mismatches < 5). Additionally, we searched for the presence of at least five pair-end reads connecting the 16S rRNA gene sequences assembled from the metagenomes (MA16S) with one or more contigs in the MAG. For all the pairs of MAGs and MA16S that satisfied these criteria, the copy number of ribosomal RNA operons per MAG was then inferred from the ratio of the coverage of the MA16S to the coverage of the MAG.

To verify that the results obtained from the assembled genomes were not biased by the use of a limited set of MAGs, the *rrn* copy number was also calculated using the whole metagenome data sets, as following. The total coverage of genomes in each dataset was estimated from the coverage of single-copy universal marker genes, identified in contigs > 500bp by utilizing Hidden Markov Models (80). Therefore, the average copy number of ribosomal RNA operons (*rrn*) per genome was calculated as the ratio of the coverage of total rRNA SSU genes (MA16S) and the total coverage of a marker gene. This calculation was repeated for a set of 13 single copy universal marker genes shorter than 450 bp (150 aa).

### Local Similarity Analysis

Local similarity analyses (LSA) (81) among the most abundant OTUs (relative abundance average > 0.05%) was used to determine associations and co-occurrence between bacterial species. Local correlations with a score lower than −0.6 and higher than 0.6, and q-value <0.01 were considered significant. The same procedure was applied to determine associations and co-occurrence between MAGs. Gephi software (82) was used to visualize and model the networks.

### Self Organizing Maps

Self Organizing Maps (SOM) were constructed using the R-package SOMbrero (83). SOM analyses were used to cluster the 60 activated sludge samples using two different criteria: I) Abundance patterns of KEGG pathway modules (84) in the whole metagenome and II) MAGs abundance. For the analysis of functional modules, genes in contigs longer than 2Kbp were predicted using the GeneMark software (85) and annotated with the GhostKOALA tool (86). Only prokaryotic modules identified as complete by the KEGG Mapper – Reconstruct Module tool (87) were used. The abundance of the pathway modules was calculated for each sample as the sum of the abundances of the genes that composed that module.

To identify the KEGG modules and MAGs showing significant differences (p < 0.05) between SOM clusters, two groups comparisons (t-test) were performed using the STAMP software (88).

### In situ growth rate determination

In-situ growth rate of bacteria in the activated sludge system was calculated using the replication index (iRep), a recently developed method which allows the direct estimation of bacterial replication rates from draft-quality genomes assembled from metagenome sequences. The index is estimated from the difference of coverage between the regions of the genome close to the single replication origin and those of the rest of the genome (27, 28). We applied the replication index (iRep) algorithm, using default parameters, for MAGs with less than 175 scaffolds per Mbp, that were more than 75% complete, had less than 5% contamination, and had a genome coverage higher than 5 in any individual sample.

## ACKNOWLEDGMENTS

This work was supported by grants from AySA-CONICET (Res. 3816/11 and 1371/15) and FONCyT (PICT 0746/15). The funders had no role in study design, data analysis, decision to publish, or preparation of the manuscript.

We are grateful to WWTP staff for help with the sampling and for sharing operation data. We also wish to thank Enrique Calderón and Omar Fioravanti for encouragement during this project.

## Competing Interests

The authors declare no competing financial interests

## SUPPLEMENTAL MATERIAL

### SUPPLEMENTAL TABLES

**Table S1:** Operational parameters and treatment performance of activated sludge wastewater treatment plant.

**Table S2:** Metagenome-assembled genomes (MAGs) information.

**Table S3:** MAGs and MA16S data used for the calculation of ribosomal RNA operon copy numbers.

### LEGENDS TO SUPPLEMENTAL FIGURES

**Fig. S1:** Nitrogen profiles over the three years of sampling. Black circles: Influent total Kjeldahl nitrogen (TKN); white circles: effluent total Kjeldahl Nitrogen; red triangles: effluent nitrate concentration; red squares: effluent nitrite concentration.

**Fig. S2:** Network of co-occurring MAGs, based on local similarity analysis (LSA). Nodes (MAGs) are colored by phylum. The size of each node is proportional to the average MAG abundance across the 60 samples. Blue and red edges represent positive and negative significant correlations (correlation > |0.6|; P-value < 0.01).

**Fig. S3:** Average relative abundance of bacterial phylum across the 60 samples, determined from the taxonomic placement of metagenome assembled genomes (MAGs), from 16S rRNA gene sequences assembled from metagenomes (MA16S) and from 16S rRNA gene amplicons (Amp16S). Bars indicated standard deviation.

**Fig. S4:** Self-organizing map (SOM) analysis. A) SOM cluster of 60 samples based on MAGs abundance and B) based on KEGG module abundance. Numbers inside boxes (clusters) correspond to samples (1 to 60). Superclusters generated using Ward’s minimum variance method are shown with colors.

**Fig. S5:** Relative abundance of KEGG orthology classes showing statistically significant differences between activated sludge samples belonging to periods of disturbance (orange bars, n = 16) and stability (green bars, n = 17). Error bars indicate within-group standard deviations. All categories had a corrected P value of <0.01 in Welch’s test. A) KEGG orthology (KO) classifications of metagenomic genes grouped into main classes. B) KEGG orthology (KO) classifications at pathway level, including genes grouped into protein families.

**Fig. S6:** MAG relative abundance showing statistically significant differences between activated sludge samples belonging to periods of disturbance (orange bars, n = 16) and stability(green bars, n = 17). Error bars indicate within-group standard deviations. Shown MAGs passed a corrected P value of <0.01 in Welch’s test.

**Fig. S7:** Heatmap of KEGG modules present in MAGs whose abundance between disturbed and stable periods were significantly different (Fig S6). MAGs were colored according to their membership to disturbed (orange) and stable (green) periods.

**Fig. S8:** Representative illustration of the relationship between MA16s and MAGs coverage used to determine the SSU operon copy number. The ID and taxonomic classification of MAGs and MA16S and estimated *rrn* copy number of 8 selected MAGs are indicated in the Figure. Lines are absolute coverage values across the 60 samples for MA16s (red lines) and MAGs (blue lines). Numbers of *rrn* copies were calculated as the average of the ratio of coverage between MA16s and MAGs. Only values above a threshold coverage = 1 (doted line) were used for calculations.

## REFERENCES

1. Hanson CA, Fuhrman JA, Horner-Devine MC, Martiny JBH. 2012. Beyond biogeographic patterns: Processes shaping the microbial landscape. Nat Rev Microbiol 10:497–506.

2. Konopka A, Lindemann S, Fredrickson J. 2015. Dynamics in microbial communities: Unraveling mechanisms to identify principles. ISME J 9:1488.

3. Nemergut DR, Schmidt SK, Fukami T, O’Neill SP, Bilinski TM, Stanish LF, Knelman JE, Darcy JL, Lynch RC, Wickey P, Ferrenberg S. 2013. Patterns and Processes of Microbial Community Assembly. Microbiol Mol Biol Rev 77:342–356.

4. Widder S, Allen RJ, Pfeiffer T, Curtis TP, Wiuf C, Sloan WT, Cordero OX, Brown SP, Momeni B, Shou W, Kettle H, Flint HJ, Haas AF, Laroche B, Kreft JU, Rainey PB, Freilich S, Schuster S, Milferstedt K, Van Der Meer JR, Grobkopf T, Huisman J, Free A, Picioreanu C, Quince C, Klapper I, Labarthe S, Smets BF, Wang H, Soyer OS, Allison SD, Chong J, Lagomarsino MC, Croze OA, Hamelin J, Harmand J, Hoyle R, Hwa TT, Jin Q, Johnson DR, de Lorenzo V, Mobilia M, Murphy B, Peaudecerf F, Prosser JI, Quinn RA, Ralser M, Smith AG, Steyer JP, Swainston N, Tarnita CE, Trably E, Warren PB, Wilmes P. 2016. Challenges in microbial ecology: Building predictive understanding of community function and dynamics. ISME J 10:2557–2568.

5. Seviour R, Nielsen PH. 2010. Microbial Ecology of Activated SludgeMicrobial Ecology. IWA Publishing, London, UK.

6. Jenkins D. 2014. Activated Sludge – 100 Years and CountingWater Intelligence Online.

7. Rittmann BE. 2006. Microbial ecology to manage processes in environmental biotechnology. Trends Biotechnol 24:261–266.

8. Saikaly PE, Stroot PG, Oertxher DB. 2005. Use of 16S rRNA gene terminal restriction fragment analysis to assess the impact of solids retention time on the bacterial diversity of activated sludge. Appl Environ Microbiol 71:5814–5822.

9. Pholchan MK, Baptista J de C, Davenport RJ, Curtis TP. 2010. Systematic study of the effect of operating variables on reactor performance and microbial diversity in laboratory-scale activated sludge reactors. Water Res 44:1341–1352.

10. Wells GF, Park HD, Eggleston B, Francis CA, Criddle CS. 2011. Fine-scale bacterial community dynamics and the taxa-time relationship within a full-scale activated sludge bioreactor. Water Res 45:5476–5488.

11. Vuono DC, Benecke J, Henkel J, Navidi WC, Cath TY, Munakata-Marr J, Spear JR, Drewes JE. 2015. Disturbance and temporal partitioning of the activated sludge metacommunity. ISME J 9:425–435.

12. Ju F, Zhang T. 2015. Bacterial assembly and temporal dynamics in activated sludge of a full-scale municipal wastewater treatment plant. ISME J 9:683–695.

13. Lee SH, Kang HJ, Park HD. 2015. Influence of influent wastewater communities on temporal variation of activated sludge communities. Water Res 73:132–144.

14. Ibarbalz FM, Figuerola ELM, Erijman L. 2013. Industrial activated sludge exhibit unique bacterial community composition at high taxonomic ranks. Water Res 47:3854–3864.

15. Gao P, Xu W, Sontag P, Li X, Xue G, Liu T, Sun W. 2016. Correlating microbial community compositions with environmental factors in activated sludge from four full-scale municipal wastewater treatment plants in Shanghai, China. Appl Microbiol Biotechnol 100:4663–4673.

16. Wang X, Hu M, Xia Y, Wen X, Ding K. 2012. Pyrosequencing analysis of bacterial diversity in 14 wastewater treatment systems in china. Appl Environ Microbiol 78:7042–7047.

17. Saunders AM, Albertsen M, Vollertsen J, Nielsen PH. 2016. The activated sludge ecosystem contains a core community of abundant organisms. ISME J 10:11–20.

18. Guo J, Ni BJ, Han X, Chen X, Bond P, Peng Y, Yuan Z. 2017. Unraveling microbial structure and diversity of activated sludge in a full-scale simultaneous nitrogen and phosphorus removal plant using metagenomic sequencing. Enzyme Microb Technol 102:16–25.

19. Griffin JS, Wells GF. 2017. Regional synchrony in full-scale activated sludge bioreactors due to deterministic microbial community assembly. ISME J 11:500–511.

20. Sheng Z, Mohammed A, Liu Y. 2016. Stability of full-scale engineered ecosystem under disturbance: Response of an activated sludge biological nutrient removal reactor to high flow rate condition. Int Biodeterior Biodegrad 109:88–95.

21. Bossier P, Verstraete W. 1996. Triggers for microbial aggregation in activated sludge? Appl Microbiol Biotechnol 45:1–6.

22. Fierer N, Breitbart M, Nulton J, Salamon P, Lozupone C, Jones R, Robeson M, Edwards RA, Felts B, Rayhawk S, Knight R, Rohwer F, Jackson RB. 2007. Metagenomic and small-subunit rRNA analyses reveal the genetic diversity of bacteria, archaea, fungi, and viruses in soil. Appl Environ Microbiol 73:7059–7066.

23. Fierer N, Bradford M a, Jackson RB. 2007. Toward an ecological classification of soil bacteria. Ecology 88:1354–64.

24. McMahon K. 2015. “Metagenomics 2.0.” Environ Microbiol Rep 7:38–39.

25. Langille MGI, Zaneveld J, Caporaso JG, McDonald D, Knights D, Reyes JA, Clemente JC, Burkepile DE, Vega Thurber RL, Knight R, Beiko RG, Huttenhower C. 2013. Predictive functional profiling of microbial communities using 16S rRNA marker gene sequences. Nat Biotechnol 31:814–821.

26. Lee ZM-PP, Bussema C, Schmidt TM. 2009. rrn DB: Documenting the number of rRNA and tRNA genes in bacteria and archaea. Nucleic Acids Res 37:D489–93.

27. Brown CT, Olm MR, Thomas BC, Banfield JF. 2016. Measurement of bacterial replication rates in microbial communities. Nat Biotechnol 34:1256–1263.

28. Olm MR, Brown CT, Brooks B, Firek B, Baker R, Burstein D, Soenjoyo K, Thomas BC, Morowitz M, Banfield JF. 2017. Identical bacterial populations colonize premature infant gut, skin, & oral microbiomes & exhibit different in situ growth rates. Genome Res 27:601–612.

29. Miller CS, Baker BJ, Thomas BC, Singer SW, Banfield JF. 2011. EMIRGE: reconstruction of full-length ribosomal genes from microbial community short read sequencing data. Genome Biol 12:R44.

30. Stoddard SF, Smith BJ, Hein R, Roller BRK, Schmidt TM. 2015. rrnDB: improved tools for interpreting rRNA gene abundance in bacteria and archaea and a new foundation for future development. Nucleic Acids Res 43:D593–D598.

31. Roller BRK, Stoddard SF, Schmidt TM. 2016. Exploiting rRNA operon copy number to investigate bacterial reproductive strategies. Nat Microbiol 1:16160.

32. Stevenson BS, Schmidt TM. 2004. Life history implications of rRNA gene copy number in Escherichia coli. Appl Environ Microbiol 70:6670–6677.

33. Roller BRK, Schmidt TM. 2015. The physiology and ecological implications of efficient growth. ISME J 9:1481–1487.

34. Wu L, Yang Y, Chen S, Jason Shi Z, Zhao M, Zhu Z, Yang S, Qu Y, Ma Q, He Z, Zhou J, He Q. 2017. Microbial functional trait of rRNA operon copy numbers increases with organic levels in anaerobic digesters. ISME J 11:2874–2878.

35. Ortiz-Álvarez R, Fierer N, de Los Ríos A, Casamayor EO, Barberán A. 2018. Consistent changes in the taxonomic structure and functional attributes of bacterial communities during primary succession. ISME J 12:1658–1667.

36. Nemergut DR, Knelman JE, Ferrenberg S, Bilinski T, Melbourne B, Jiang L, Violle C, Darcy JL, Prest T, Schmidt SK, Townsend AR. 2016. Decreases in average bacterial community rRNA operon copy number during succession. ISME J 10:1147–1156.

37. Niederdorfer R, Besemer K, Battin TJ, Peter H. 2017. Ecological strategies and metabolic trade-offs of complex environmental biofilms. npj Biofilms Microbiomes 3.

38. Pianka ER. 1970. On r- and K-Selection. Am Nat 104:592–597.

39. Ho A, Di Lonardo DP, Bodelier PLE. 2017. Revisiting life strategy concepts in environmental microbial ecology. FEMS Microbiol Ecol 93:fix006.

40. De Vries FT, Shade A. 2013. Controls on soil microbial community stability under climate change. Front Microbiol 4:265.

41. Luo H, Csuros M, Hughes A, Moran M. 2013. Evolution of divergent life history strategies in marine alphaproteobacteria. MBio 4:e00373–13.

42. Levin BR, McCall IC, Perrot V, Weiss H, Ovesepian A, Baquero F. 2017. A numbers game: Ribosome densities, bacterial growth, and antibiotic-mediated stasis and death. MBio 8:e02253–16.

43. Korem T, Zeevi D, Suez J, Weinberger A, Avnit-Sagi T, Pompan-Lotan M, Matot E, Jona G, Harmelin A, Cohen N, Sirota-Madi A, Thaiss CA, Pevsner-Fischer M, Sorek R, Xavier RJ, Elinav E, Segal E. 2015. Growth dynamics of gut microbiota in health and disease inferred from single metagenomic samples. Science 349:1101–1106.

44. Hu P, Dubinsky EA, Probst AJ, Wang J, Sieber CMK, Tom LM, Gardinali PR, Banfield JF, Atlas RM, Andersen GL. 2017. Simulation of Deepwater Horizon oil plume reveals substrate specialization within a complex community of hydrocarbon degraders. Proc Natl Acad Sci 114:7432–7437.

45. Costello EK, Sun CL, Carlisle EM, Morowitz MJ, Banfield JF, Relman DA. 2017. Candidatus Mycoplasma girerdii replicates, diversifies, and co-occurs with Trichomonas vaginalis in the oral cavity of a premature infant. Sci Rep 7:3764.

46. Probst AJ, Ladd B, Jarett JK, Geller-Mcgrath DE, Sieber CMK, Emerson JB, Anantharaman K, Thomas BC, Malmstrom RR, Stieglmeier M, Klingl A, Woyke T, Ryan MC, Banfield JF. 2018. Differential depth distribution of microbial function and putative symbionts through sediment-hosted aquifers in the deep terrestrial subsurface. Nat Microbiol 3:328.

47. Manhart M, Adkar B V., Shakhnovich EI. 2018. Trade-offs between microbial growth phases lead to frequency-dependent and non-transitive selection. Proc R Soc B Biol Sci 285:2459.

48. Condon C, Liveris D, Squires C, Schwartz I, Squires CL. 1995. rRNA operon multiplicity in Escherichia coli and the physiological implications of rrn inactivation. J Bacteriol 177:4152–4156.

49. Gyorfy Z, Draskovits G, Vernyik V, Blattner FF, Gaal T, Posfai G. 2015. Engineered ribosomal RNA operon copy-number variants of E. coli reveal the evolutionary trade-offs shaping rRNA operon number. Nucleic Acids Res 43:1783–1794.

50. Adkar B V., Manhart M, Bhattacharyya S, Tian J, Musharbash M, Shakhnovich EI. 2017. Optimization of lag phase shapes the evolution of a bacterial enzyme. Nat Ecol Evol 1:149.

51. Swinnen IAM, Bernaerts K, Dens EJJ, Geeraerd AH, Van Impe JF. 2004. Predictive modelling of the microbial lag phase: A review. Int J Food Microbiol 94:137–159.

52. Lauro FM, McDougald D, Thomas T, Williams TJ, Egan S, Rice S, DeMaere MZ, Ting L, Ertan H, Johnson J, Ferriera S, Lapidus A, Anderson I, Kyrpides N, Munk AC, Detter C, Han CS, Brown M V., Robb FT, Kjelleberg S, Cavicchioli R. 2009. The genomic basis of trophic strategy in marine bacteria. Proc Natl Acad Sci 106:15527–15533.

53. Haggerty JM, Dinsdale EA. 2017. Distinct biogeographical patterns of marine bacterial taxonomy and functional genes. Glob Ecol Biogeogr 26:177–190.

54. Korem Kohanim Y, Levi D, Jona G, Towbin BD, Bren A, Alon U. 2018. A Bacterial Growth Law out of Steady State. Cell Rep 23:2891–2900.

55. Mori M, Schink S, Erickson DW, Gerland U, Hwa T. 2017. Quantifying the benefit of a proteome reserve in fluctuating environments. Nat Commun.

56. Frolov EN, Belousova E V., Lavrinenko KS, Dubinina GA, Grabovich MY. 2013. Capacity of Azospirillum thiophilum for lithotrophic growth coupled to oxidation of reduced sulfur compounds. Microbiology 82:271–279.

57. Parkinson JS, Kofoid EC. 1992. Communication Modules in Bacterial Signaling Proteins. Annu Rev Genet 26:71–112.

58. Shade A, Peter H, Allison SD, Baho DL, Berga M, Bürgmann H, Huber DH, Langenheder S, Lennon JT, Martiny JBH, Matulich KL, Schmidt TM, Handelsman J. 2012. Fundamentals of microbial community resistance and resilience. Front Microbiol 3:417.

59. Harley CDG, Paine RT. 2009. Contingencies and compounded rare perturbations dictate sudden distributional shifts during periods of gradual climate change. Proc Natl Acad Sci 106:11172–11176.

60. Plante CJ. 2017. Defining Disturbance for Microbial Ecology. Microb Ecol 74:259–263.

61. Henze M, Grady CPL, Gujer W, Marais G, Matsuo T. 1987. Activated sludge model No. 1. Task group on mathematical modelling for design and operation of biological wastewater treatmentScientific and Technical Rep.

62. Akarsubasi AT, Eyice O, Miskin I, Head IM, Curtis TP. 2009. Effect of sludge age on the bacterial diversity of bench scale sequencing batch reactors. Environ Sci Technol 43:2950–2956.

63. Schultz JR, Hegg BA, Rakness KL. 1982. Realistic Sludge Production for Activated Sludge Plants without Primary Clarifiers. J Water Pollut Control Fed 54:1355–1360.

64. Kappeler J, Gujer W. 1992. Estimation of kinetic parameters of heterotrophic biomass under aerobic conditions and characterization of wastewater for activated sludge modelling, p. 125–139. *In* Water Science and Technology.

65. Schimel J, Balser TC, Wallenstein M. 2007. Microbial stress-response physiology and its implications for ecosystem function. Ecology 88:1386–1394.

66. Guénon R, Gros R. 2013. Frequent-wildfires with shortened time-since-fire affect soil microbial functional stability to drying and rewetting events. Soil Biol Biochem 57:663–674.

67. Lennon JT, Aanderud ZT, Lehmkuhl BK, Schoolmaster DR. 2012. Mapping the niche space of soil microorganisms using taxonomy and traits. Ecology 93:1867–1879.

68. Ibarbalz FM, Perez MV, Figuerola ELM, Erijman L. 2014. The bias associated with amplicon sequencing does not affect the quantitative assessment of bacterial community dynamics. PLoS One 9:e99722.

69. Herlemann DPR, Labrenz M, Jürgens K, Bertilsson S, Waniek JJ, Andersson AF. 2011. Transitions in bacterial communities along the 2000 km salinity gradient of the Baltic Sea. ISME J 5:1571–1579.

70. Bolger AM, Lohse M, Usadel B. 2014. Trimmomatic: A flexible trimmer for Illumina sequence data. Bioinformatics 30:2114–2120.

71. Edgar RC. 2010. Search and clustering orders of magnitude faster than BLAST. Bioinformatics 26:2460–2461.

72. Quast C, Pruesse E, Yilmaz P, Gerken J, Schweer T, Yarza P, Peplies J, Glöckner FO. 2013. The SILVA ribosomal RNA gene database project: improved data processing and web-based tools. Nucleic Acids Res 41:D590–D596.

73. Li D, Liu CM, Luo R, Sadakane K, Lam TW. 2015. MEGAHIT: An ultra-fast single-node solution for large and complex metagenomics assembly via succinct de Bruijn graph. Bioinformatics 31:1674–1676.

74. Kang DD, Froula J, Egan R, Wang Z. 2015. MetaBAT, an efficient tool for accurately reconstructing single genomes from complex microbial communities. PeerJ 3:e1165.

75. Shannon P, Markiel A, Ozier O, Baliga NS, Wang JT, Ramage D, Amin N, Schwikowski B, Ideker T. 2003. Cytoscape: a software environment for integrated models of biomolecular interaction networks. Genome Res 13:2498–2504.

76. Parks DH, Imelfort M, Skennerton CT, Hugenholtz P, Tyson GW. 2015. CheckM: Assessing the quality of microbial genomes recovered from isolates, single cells, and metagenomes. Genome Res 25:1043–1055.

77. Langmead B, Salzberg SL. 2012. Fast gapped-read alignment with Bowtie 2. Nat Methods 9:357–359.

78. Parks DH, Chuvochina M, Waite DW, Rinke C, Skarshewski A, Chaumeil PA, Hugenholtz P. 2018. A standardized bacterial taxonomy based on genome phylogeny substantially revises the tree of life. Nat Biotechnol 36:996.

79. Bengtsson-Palme J, Hartmann M, Eriksson KM, Pal C, Thorell K, Larsson DGJ, Nilsson RH. 2015. metaxa2: improved identification and taxonomic classification of small and large subunit rRNA in metagenomic data. Mol Ecol Resour 15:1403–1414.

80. Wu M, Scott AJ. 2012. Phylogenomic analysis of bacterial and archaeal sequences with AMPHORA2. Bioinformatics 28:1033–1034.

81. Xia LC, Steele JA, Cram JA, Cardon ZG, Simmons SL, Vallino JJ, Fuhrman JA, Sun F. 2011. Extended local similarity analysis (eLSA) of microbial community and other time series data with replicates. BMC Syst Biol 5.

82. Bastian M, Heymann S, Jacomy M. 2009. Gephi: An Open Source Software for Exploring and Manipulating Networks. Third Int AAAI Conf Weblogs Soc Media 8:361–362.

83. Boelaert J, Bendhaiba L, Olteanu M, Villa-Vialaneix N. 2014. SOMbrero: An R Package for Numeric and Non-numeric Self-Organizing Maps, p. 219–228. In Advances in Intelligent Systems and Computing.

84. Kanehisa M. 2000. KEGG: Kyoto Encyclopedia of Genes and Genomes. Nucleic Acids Res 28:27–30.

85. Besemer J, Borodovsky M. 2005. GeneMark: Web software for gene finding in prokaryotes, eukaryotes and viruses. Nucleic Acids Res 33.

86. Kanehisa M, Sato Y, Morishima K. 2016. BlastKOALA and GhostKOALA: KEGG Tools for Functional Characterization of Genome and Metagenome Sequences. J Mol Biol 428:726–731.

87. Kanehisa M, Goto S, Sato Y, Furumichi M, Tanabe M. 2012. KEGG for integration and interpretation of large-scale molecular data sets. Nucleic Acids Res 40.

88. Parks DH, Tyson GW, Hugenholtz P, Beiko RG. 2014. STAMP: Statistical analysis of taxonomic and functional profiles. Bioinformatics 30:3123–3124.

89. Water Environment Federation (WEF). 2008. Operation of Municipal Wastewater Treatment Plants. McGraw-Hill, Chicago, USA.

